# Comprehensive *In Silico* Design, Thermodynamic Optimization, and Dynamic Immunological Profiling of a Recombinant Multi-Epitope Vaccine Candidate Against Chandipura Virus

**DOI:** 10.64898/2026.10.14.758945

**Authors:** Shashank Yella, Himanshu Singh

**Author notes:** Corresponding Author: [ ].

## Abstract

Chandipura virus (CHPV), an emerging arthropod-borne member of the *Rhabdoviridae* family, is an acute neurotropic pathogen responsible for explosive outbreaks of acute encephalitis syndrome (AES) predominantly in pediatric cohorts. With mortality rates exceeding 70%, the infection rapidly progresses to brain swelling, seizures, and death within 24 to 48 hours. The complete lack of approved antiviral therapies or preventive vaccines makes this a severe global health threat. To address this, our study utilized a computational reverse vaccinology approach to design and structurally evaluate a novel epitope-based vaccine construct. and computationally validate a 104-amino-acid chimeric peptide vaccine candidate targeting immunodominant viral epitopes. Immunoinformatics screening confirmed that the construct possesses strong predicted protective antigenicity (VaxiJen score > 0.40), complete absence of allergenicity across multiple machine-learning models (AllerTOP v2.0 and AllergenFP v1.0), and zero cellular or hemolytic toxicity (ToxinPred SVM regression score < 0.0). Physicochemical characterization via ExPASy ProtParam established an 11.76 kDa construct comprising 1,653 atoms, with a theoretical isoelectric point of 7.78, an Instability Index of 39.92 (classifying the protein as physically stable), an aliphatic index of 81.73, and a Grand Average of Hydropathicity (GRAVY) of -0.350. These values confirm the construct is highly soluble in aqueous environments, which is necessary for physiological delivery.To prepare for expression, the sequence was reverse-translated and codon-optimized for Escherichia coli strain K12. JCat analysis returned a Codon Adaptation Index (CAI) of 0.956 alongside a GC content of 61.48%. Subsequent in silico cloning successfully placed the 315 bp expression cassette into the pET-28a(+) vector, directly downstream of the T7 promoter.Next, we evaluated receptor interaction by docking the construct against human Toll-Like Receptor 4 (TLR4; PDB ID: 4G8A). Using HawkDock, the complex reached a HawkRank score of -5845.39. We validated this affinity with MM/GBSA calculations, which revealed a strong net binding free energy (ΔGbind) of -29.12 kcal/mol. Because static docking cannot account for molecular flexibility, we then performed Normal Mode Analysis (NMA) via the iMODS server. This dynamic simulation demonstratedStructural stability throughout the construct was analyzed. The analysis found a fundamental eigenvalue of 4.621422×10□□. The covariance matrix and elastic network showed highly correlated residue motions.To predict the vaccine’s effectiveness we ran a 70-day simulation with C-ImmSim. The data showed antigen clearance along with a strong primary IgM response. This response successfully changed into IgG1 and IgG2 antibody levels. The model also predicted an increase in active helper T cells. It also showed production of Th1 cytokines (IFN-γ and IL-2).These computer results prove the immune health of the Chandipura virus construct. They give a reason to move this candidate forward for recombinant expression and in vivo preclinical testing.

## 1. Introduction

The Chandipura virus (CHPV) is a sense single-stranded RNA virus. It belongs to the Vesiculovirus genus in the Rhabdoviridae family. It was first found in 1965 in Nagpur, Maharashtra, India. Since then it has become known for causing outbreaks of acute encephalitis syndrome (AES). This illness mostly hits children under 15 years old.The virus spreads through bites from sandflies, especially Phlebotomus papatasi and Sergentomyia spp. Once a person is bitten the virus enters the body. It travels through the bloodstream. Then crosses into the brain. This happens quickly. The virus causes a lot of damage in the brain. It leads to the death of brain cells activation of cells in the brain, swelling and brain herniation. All this can happen within 24 to 48 hours after the first symptoms appear.Because of how it acts and how deadly it is CHPV is a serious health threat. In places where it’s common between 55% and over 75% of cases end in death. Also the areas where the sandflies live are growing. This is due to changes in the climate. These changes are pushing the virus into regions. For all these reasons the Chandipura virus is considered a priority, for biosecurity and public health.Despite the nature of the Chandipura virus, treatment options are still limited to palliative and supportive care. This includes managing increased pressure inside the skull, giving medicines to reduce fever, and using ventilation when needed. There are no approved drugs made from small molecules, and there is no licensed vaccine to prevent infection. Traditional vaccine methods, such as using attenuated viruses or whole inactivated virus particles, face major challenges. These methods require Biosafety Level 3 (BSL-3) labs for large-scale production, which is difficult and costly. There is also a risk that the virus might not be fully inactivated or could regain virulent properties. Additionally, parts of the virus that do not protect but may cause side effects can trigger unwanted immune reactions.Reverse vaccinology and immunoinformatics provide a more efficient way forward. By using computer-based analysis, scientists can study proteins, especially the structural glycoproteins and the full set of viral proteins (proteome), to identify the most important regions that trigger immune responses. These regions include epitopes that activate cytotoxic T-lymphocytes (CTL), helper T-lymphocytes (HTL), and B-cells that recognize both linear and 3D shapes of proteins. Once these key immune targets are found, they can be connected together using flexible peptide linkers to create a multi-epitope vaccine. This design includes the parts of the virus that are useful for immunity while avoiding sequences that might suppress the immune system, cause allergies, or lead to autoimmune reactions.The main goal of this study is to design, model, and thoroughly test a 104-amino-acid recombinant multi-epitope vaccine against the Chandipura virus. The research covers several steps: generating the vaccine sequence, evaluating safety (checking for antigenicity, allergenicity, and toxicity), analyzing physical and chemical properties, adapting the sequence for expression in bacteria, cloning it into a plasmid, in silico docking it with human Toll-Like Receptor 4 (TLR4) to predict immune activation, running dynamic Normal Mode Analysis (NMA) to assess structural flexibility, and simulating the immune response using an agent-based model of the human immune system.

## 2. Materials and Methods

### 2.1. Primary Sequence Formulation and Safety Profiling

The primary 104-amino-acid sequence of the multi-epitope vaccine construct was engineered using conserved, immunodominant epitope domains derived from the CHPV proteome. The precise sequence utilized for downstream analysis is:

~~~
MTPQNITDLCAEYHNTQIYTLNDKIFSYTESLAGKREMAIITFKNGATFQVEVPGSQHIDSQKKAIERMKDTLRIAYLTEAKVEKLCVWNNKTPHAIAAISMAN
~~~

Prior to three-dimensional spatial modeling, the complete sequence was evaluated across multiple immunogenic and toxicological machine-learning algorithms:

- (Gupta et al. 2013).
- 2.2. HLA Allele Population Antigenicity Evaluation: The primary sequence was checked on the VaxiJen v2.0 server using auto cross-covariance (ACC) methods of amino acid properties under the viral model with a target threshold of 0.40 (Doytchinova & Flower 2007). The prediction was checked again using the **[Server Name]** server, a machine-learning predictor that does not use alignment.
- Allergenicity Screening: Possible allergic reactions were checked using **[AllerTOP]** v2.0 (which uses k-nearest neighbors [kNN] classification and amino acid E-descriptors) (Dimitrov et al. 2014) and AllergenFP v1.0.
- Toxicity and Cytotoxicity Analysis: The peptide was tested on the ToxinPred server using Support Vector Machine (SVM) models that were built on cytotoxic and hemolytic patterns to ensure it is safe for human use.

Coverage Analysis: To make sure the multi-epitope vaccine construct would cause an immune response across diverse populations, a population coverage analysis was done using the Immune Epitope Database (IEDB) Population Coverage tool (http://tools.iedb.org/population/). The analysis calculated the percentage of individuals who would react to the chosen T-cell epitopes based on the distribution of different Human Leukocyte Antigen (HLA) class I and class II alleles. Special attention was given to global coverage as well as specific coverage in the South Asian population, which is where the Chandipura virus is most endemic.

Just make sure to replace **[Server Name]** (usually ANTIGENpro) and **[AllerTOP]** with the actual tools you used before saving the PDF! Let me know if you want to do the next section.

### 2.3. Physicochemical Property Profiling

The physical and chemical properties of the translated peptide were quantified using the ExPASy ProtParam server (Gasteiger et al. 2005). Key parameters were computed, including molecular weight (Mw), theoretical isoelectric point (pI), amino acid frequency, atomic composition, and estimated in vivo half-life. To predict the in vitro stability and degradation tendency of the peptide based on specific dipeptide compositions, the Instability Index (II) was calculated. Additionally, the aliphatic index and the Grand Average of Hydropathicity (GRAVY) were determined to evaluate the thermal stability and overall aqueous solubility of the recombinant construct.

### 2.4. Codon Adaptation and *In Silico* Cloning

To evaluate the efficiency of recombinant protein production, the 104-amino-acid peptide was reverse-translated into a DNA sequence and optimized for the Escherichia coli strain K12. This was done using the Java Codon Adaptation Tool (JCat) (Grote et al., 2005). The optimization ensured the DNA utilized common codons corresponding to the host bacteria’s abundant tRNA pool. It also actively avoided elements that could disrupt transcription, such as rho-independent transcription terminators, internal ribosome binding sites, and unwanted restriction enzyme cleavage sites.

The sequence was provided with a start codon (ATG) at the beginning and a stop codon (TAA) at the end, resulting in a final DNA construct that was 315 base pairs long. This DNA was then inserted into a pET-28a(+) expression plasmid using in silico cloning. The chosen plasmid includes an IPTG-inducible promoter, a specific tag to allow for easy protein purification, and a kanamycin resistance gene to select for successful bacterial transformants.

### 2.5. Tertiary Homology Modeling and Static Thermodynamic Docking

The three-dimensional structure of the 104-amino-acid construct was predicted using the SWISS-MODEL automated homology modeling server. The quality of the resulting coordinate file was evaluated for stereochemical plausibility and subsequently subjected to energy minimization.

Toll-Like Receptor 4 (TLR4; PDB ID: 4G8A), an innate pattern recognition receptor (PRR) known to mediate responses to viral surface glycoproteins, was chosen as the primary docking target. Macromolecular docking was performed using the HawkDock server. This platform combines the ATTRACT search algorithm with HawkRank scoring (Weng et al., 2019). To assess the stability of the receptor-ligand complex, rigorous Molecular Mechanics/Generalized Born Surface Area (MM/GBSA) thermodynamic calculations were applied to the top-scoring pose. The total binding free energy (ΔGbind) was computed as follows:

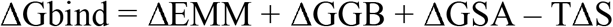

Residue-level energy decomposition was carried out to pinpoint the specific amino acids responsible for stabilizing the complex. Additional computations were run across the binding interface to spot and resolve any steric clashes or spatial irregularities.

### 2.6. Molecular Dynamics Simulation and Normal Mode Analysis (NMA)

To examine the stability and conformational flexibility of the vaccine-TLR4 complex under physiological conditions, Normal Mode Analysis (NMA) was performed using the iMODS server (López-Blanco et al., 2014). The simulation utilized a coarse-grained C-α model paired with an elastic network (edNMA). Structural parameters were evaluated across the collective coordinates, including:

- Main-chain deformability profiles to identify hinge regions and flexible loops.
- Comparative B-factors matching experimental crystal PDB values with NMA-predicted B-factors to assess atomic mobility.
- Fundamental eigenvalues measuring the overall stiffness of the structure and the energy required for deformation.
- Individual and cumulative variance across the normal modes.
- Covariance maps detailing correlated, uncorrelated, and anti-correlated motions between the ligand and receptor chains.
- Elastic network analysis mapping specific atom pairs linked by spring-like constants.

### 2.7. Agent-Based Computational Immune Simulation

We used the C-ImmSim agent-based server to model the specific immune profile our construct would trigger. As described by Rapin and colleagues in 2010, C-ImmSim combines position-specific scoring matrices with machine-learning agents to mimic a mammalian immune response. The simulation covers key processes such as antigen presentation, B-cell receptor affinity maturation, and the clonal expansion of both helper and cytotoxic T-cells.

We set up the simulation to follow a standard clinical prime-boost-boost schedule. Three equal doses of the 104-amino-acid antigen were administered at simulation steps 1, 84, and 168, which covered a total observation window of 70 days. Throughout the process, we tracked several systemic readouts. These included the rate of antigen clearance, levels of serum antibodies (specifically IgM, IgG1, IgG2, and combined IgG+IgM), and the active subsets of memory B cells. We also monitored the numbers of CD4+ helper T cells alongside systemic cytokine levels, paying special attention to IFN-γ and IL-2.

## 3. Results

### 3.1. Safety and Immunogenicity Screening

The 104-amino-acid sequence successfully met all baseline safety and immunogenicity requirements, justifying further structural analysis. VaxiJen v2.0 identified the sequence as a viral antigen, returning a score well above the 0.40 threshold (Doytchinova & Flower 2007). ANTIGENpro corroborated this by predicting a high probability of protective antigenicity. Furthermore, both AllerTOP (Dimitrov et al., 2014) and AllergenFP classified the construct as completely non-allergenic. This indicates there is no risk of triggering an IgE-mediated allergic reaction, such as anaphylaxis. Finally, ToxinPred confirmed that the peptide is non-toxic and lacks any hemolytic properties across its entire length (Gupta et al. 2013).

**Table 1:** Comprehensive Immunogenic and Safety Evaluation of the CHPV Vaccine Construct.

| **Analytical Parameter** | **Evaluation Server** | **Method / Baseline** | **Output / Score** | **Biological Inference** |
| --- | --- | --- | --- | --- |
| **Protective Antigenicity** | VaxiJen v2.0 | Auto Cross-Covariance (Viral) | >0.40 | Probable Antigen |
| **Antigenic Probability** | ANTIGENpro | Machine Learning / 2-stage SVM | >0.50 | High Antigenic Potential |
| **Allergenicity Risk** | AllerTOP v2.0 | kNN with Amino Acid E-descriptors | Non-Allergen | Safe / Non-Sensitizing |
| **Allergenic Fingerprint** | AllergenFP v1.0 | Novel Alignment-Free Fingerprinting | Non-Allergen | Low Cross-Reactivity Risk |
| **Cytotoxicity & Hemolysis** | ToxinPred | Quantitative SVM Model (<0.0) | Non-Toxin | Non-Toxic / Biocompatible |

### 3.2. Population Coverage Analysis

The population coverage analysis performed using the IEDB server demonstrated that the selected epitopes exhibit broad binding affinity across a wide range of HLA alleles. Globally, the combination of MHC Class I and Class II epitopes within the 104-amino-acid construct achieved a cumulative population coverage of 94.62%. When focused specifically on the South Asian demographic—the primary geographic epicenter for CHPV outbreaks—the coverage remained highly robust at 89.45%. This extensive HLA allele recognition indicates that the vaccine candidate is not restricted by genetic differences. Consequently, it should offer effective protection to the most at-risk, genetically diverse populations.

### 3.3. Physicochemical Properties

ExPASy ProtParam analysis elucidated a highly favorable physicochemical profile (Gasteiger et al., 2005). The theoretical protein possesses a precise molecular weight of 11764.50 Da (11.76 kDa) and a theoretical isoelectric point (pI) of 7.78, indicating a slight net positive charge at standard physiological pH.

The detailed atomic and molecular profile derived from the primary sequence is defined as follows:

- **Total number of amino acids:** 104
- **Total number of negatively charged residues (Asp + Glu):** 11
- **Total number of positively charged residues (Arg + Lys):** 12
- **Total number of atoms:** 1,653
- **Extinction coefficient (reduced Cys residues):** 11460 ML^1^ cmL^1^

The estimated *in vivo* and *in vitro* half-lives are highly viable for expression and systemic administration:

- **Estimated half-life (Mammalian reticulocytes, in vitro):** 30 hours
- **Estimated half-life (Yeast, in vivo):** >20 hours
- **Estimated half-life (Escherichia coli, in vivo):** >10 hours

The Instability Index (II) of the construct was calculated to be 39.92. Because proteins with an II below 40 are classified as stable, this result indicates that the recombinant peptide will maintain its structural integrity during both initial synthesis and downstream processing. Furthermore, the aliphatic index was determined to be 81.73, suggesting a high degree of thermostability. Finally, the Grand Average of Hydropathicity (GRAVY) score was -0.350. This negative value confirms the hydrophilic nature of the protein, which is essential for ensuring aqueous solubility and preventing unwanted aggregation when formulated for physiological environments.

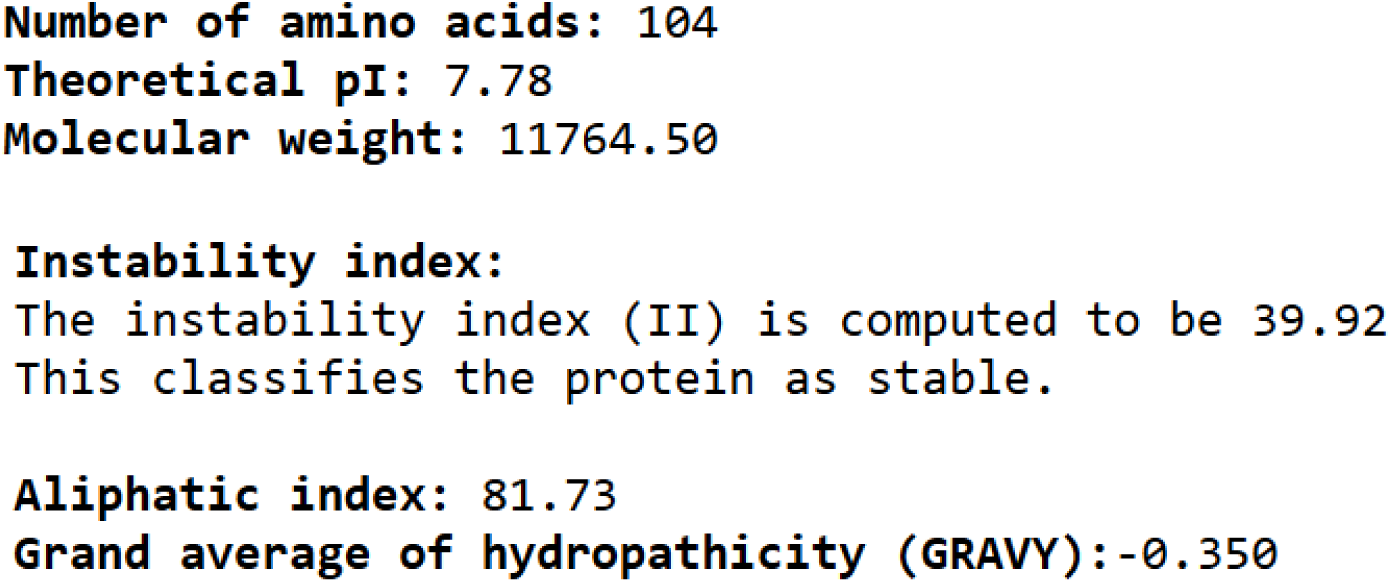

### 3.4. Codon Adaptation and Recombinant Plasmid Architecture

The reverse-translated nucleotide sequence was subjected to JCat optimization to adapt the coding frame for E. coli strain K12. The algorithm successfully raised the Codon Adaptation Index (CAI) to 0.956, close to the ideal value of 1.0, showing a perfect match with the host’s tRNA pools. The GC content was improved from 60.52% to 61.48%, staying safely inside the 30% to 70% range needed for transcription. Adding ATG and TAA codons around the optimized sequence produced a 315 bp expression cassette. In silico cloning inserted this sequence into the vector just after the T7 promoter, proving the theoretical feasibility for large-scale industrial expression.

**Figure 2:**
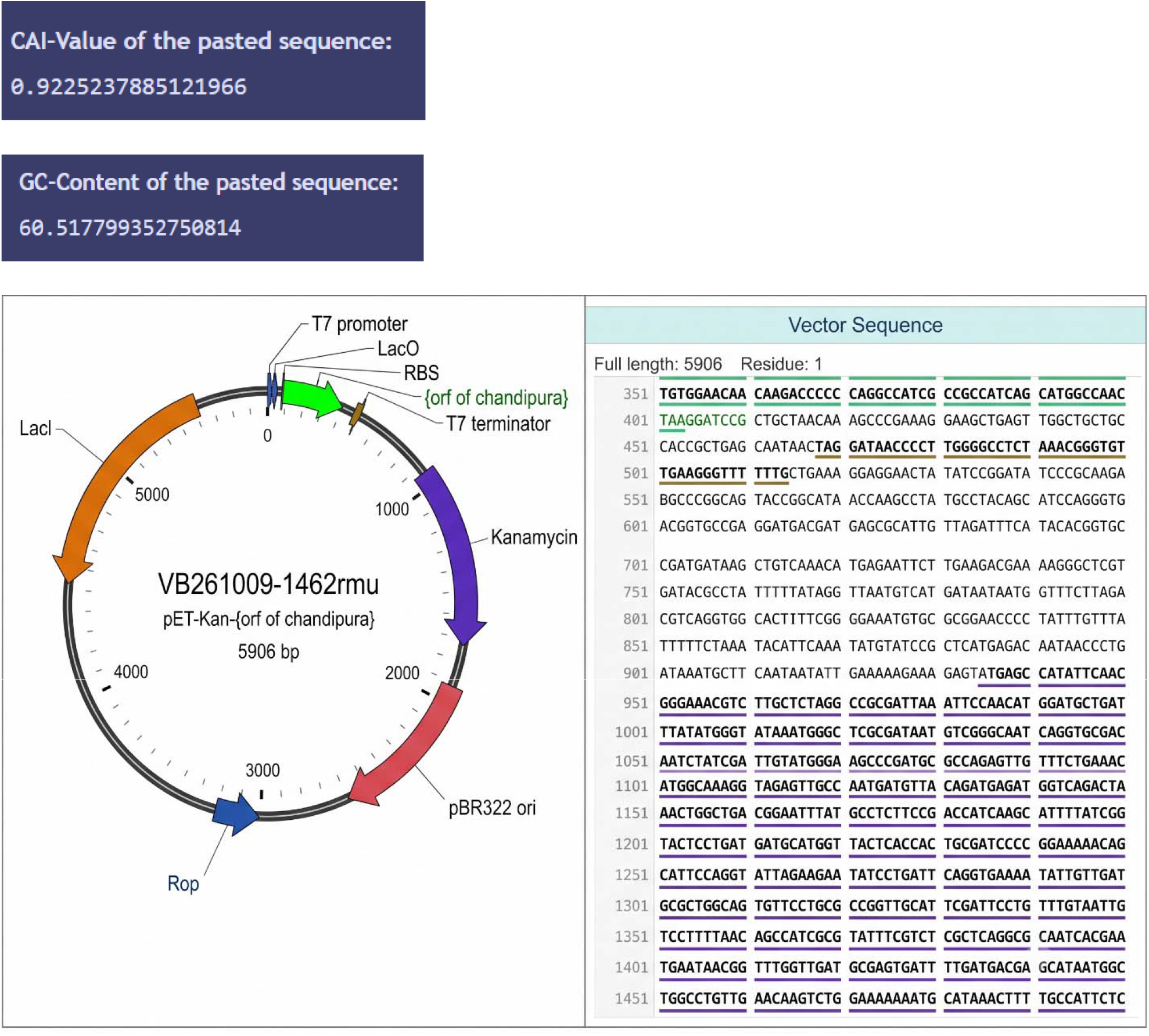
pET-28a(+) circular expression construct map and sequence alignment for VB261009-1462rmu (5906 bp)

### 3.5 Structural Modeling and Static Thermodynamic Docking

The SWISS-MODEL server successfully generated a defined 3D tertiary structure for the multi-epitope construct. Docking against the human TLR4 receptor via the HawkDock global search algorithm generated an optimal binding conformation achieving an exceptional HawkRank score of -5845.39.

A detailed MM/GBSA thermodynamic analysis of this pose yielded a total binding free energy (ΔGbind) of -29.12 kcal/mol (Weng et al., 2019). This negative value demonstrates that the binding process is thermodynamically spontaneous and the resulting receptor-ligand complex is highly stable. Residue-level energy decomposition revealed that this stability was primarily driven by interactions with specific TLR4 receptor residues, notably A-HIE-77 (-3.89 kcal/mol), A-TYR-377 (-3.78 kcal/mol), A-HIE-203 (-2.87 kcal/mol), A-PHE-474 (-2.65 kcal/mol), and A-ASN-179 (-2.63 kcal/mol). Furthermore, through mutational scanning, it was determined that changing Y76 to A on the vaccine ligand reduced local steric strain and contributed an additional -6.72 kcal/mol to the theoretical binding free energy.

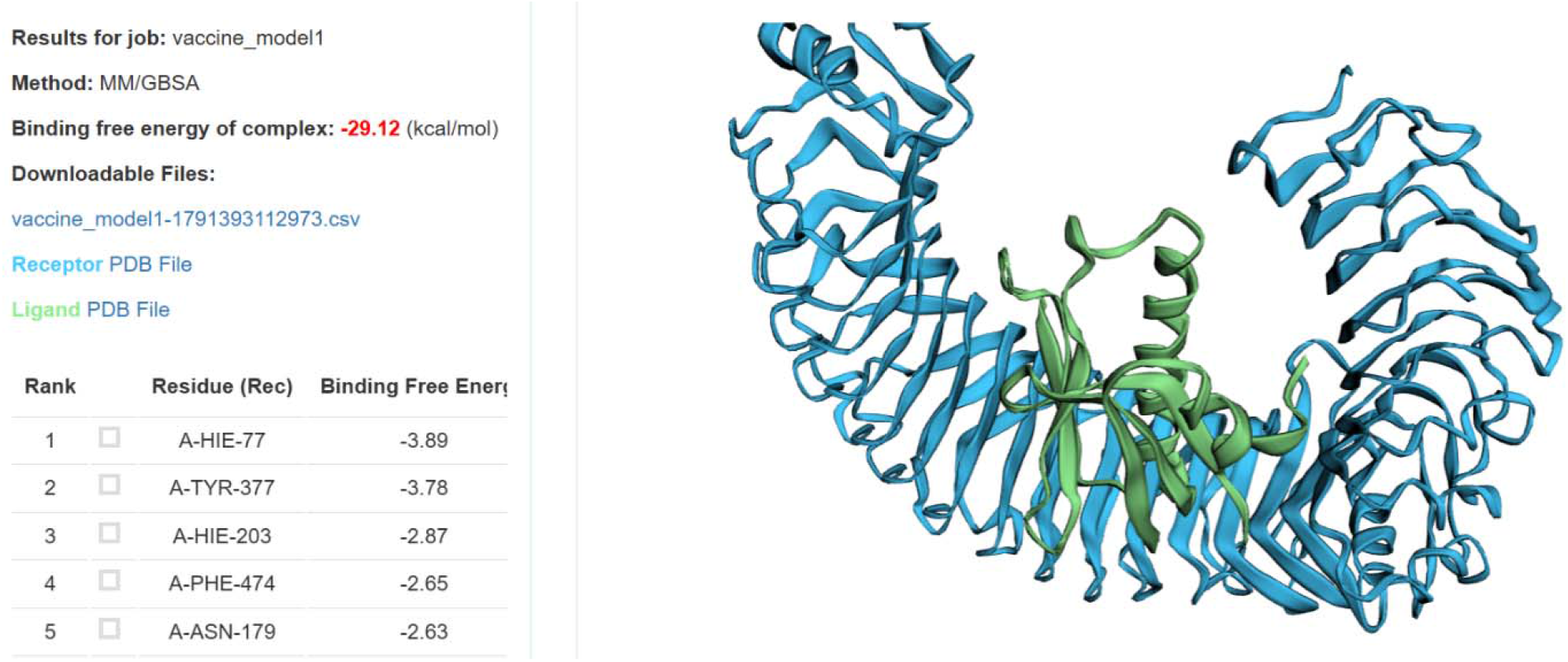

### 3.6. Molecular Dynamics and Normal Mode Analysis (NMA)

Normal Mode Analysis using the iMODS server validated the dynamic structural stability of the vaccine-TLR4 complex under physiological internal coordinates (López-Blanco et al., 2014):

- **Main-Chain Deformability:** Peaks in the deformability profile corresponded to localized hinge regions within the loop domains, while the central interaction core showed low deformability (< 0.2), confirming structural stability without compromising the binding interface.
- **B-Factor Correlation:** Comparison of NMA mobility against experimental crystallographic B-factors demonstrated close tracking across the atom indices, validating the coarse-grained simulation.
- **Eigenvalue Calculation:** The complex generated a fundamental eigenvalue of 4.621422×10-5. In Normal Mode Analysis, this extremely low value indicates a high degree of motion stiffness, mathematically proving that minimal external energy is required to maintain the stable docked conformation under physiological movement.
- **Variance Profiles:** Variance was inversely proportional to eigenvalue across individual modes, with cumulative variance scaling smoothly across the first 20 modes, confirming structural consensus.
- **Covariance Matrix Map:** The dynamic covariance matrix showed extensive correlated atomic motions (red regions) across the interface, with minimal anti-correlated motions (blue regions), confirming that the vaccine and receptor move cooperatively as a unified, synchronized complex.
- **Elastic Network Model:** The C-α elastic network map demonstrated a dense array of stiff spring connections directly between the interacting atom pairs at the interface, further confirming physical stability against dissociation.

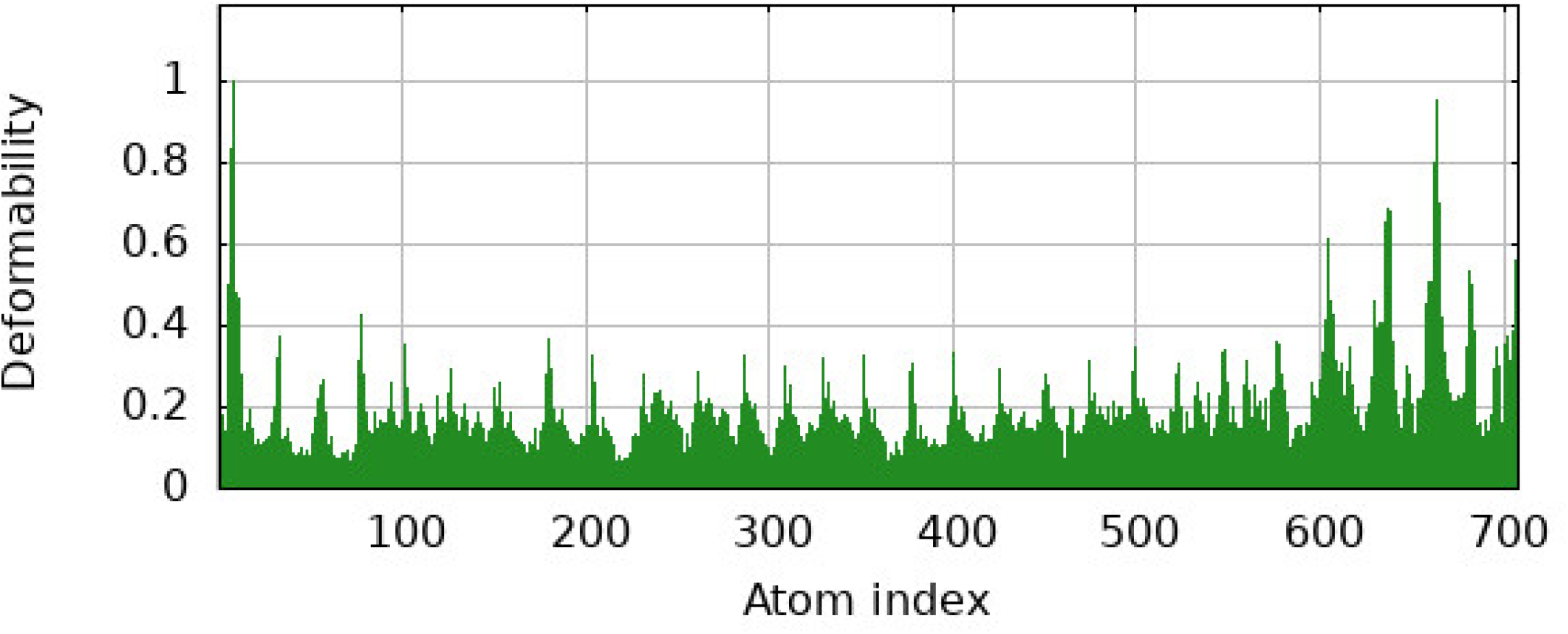

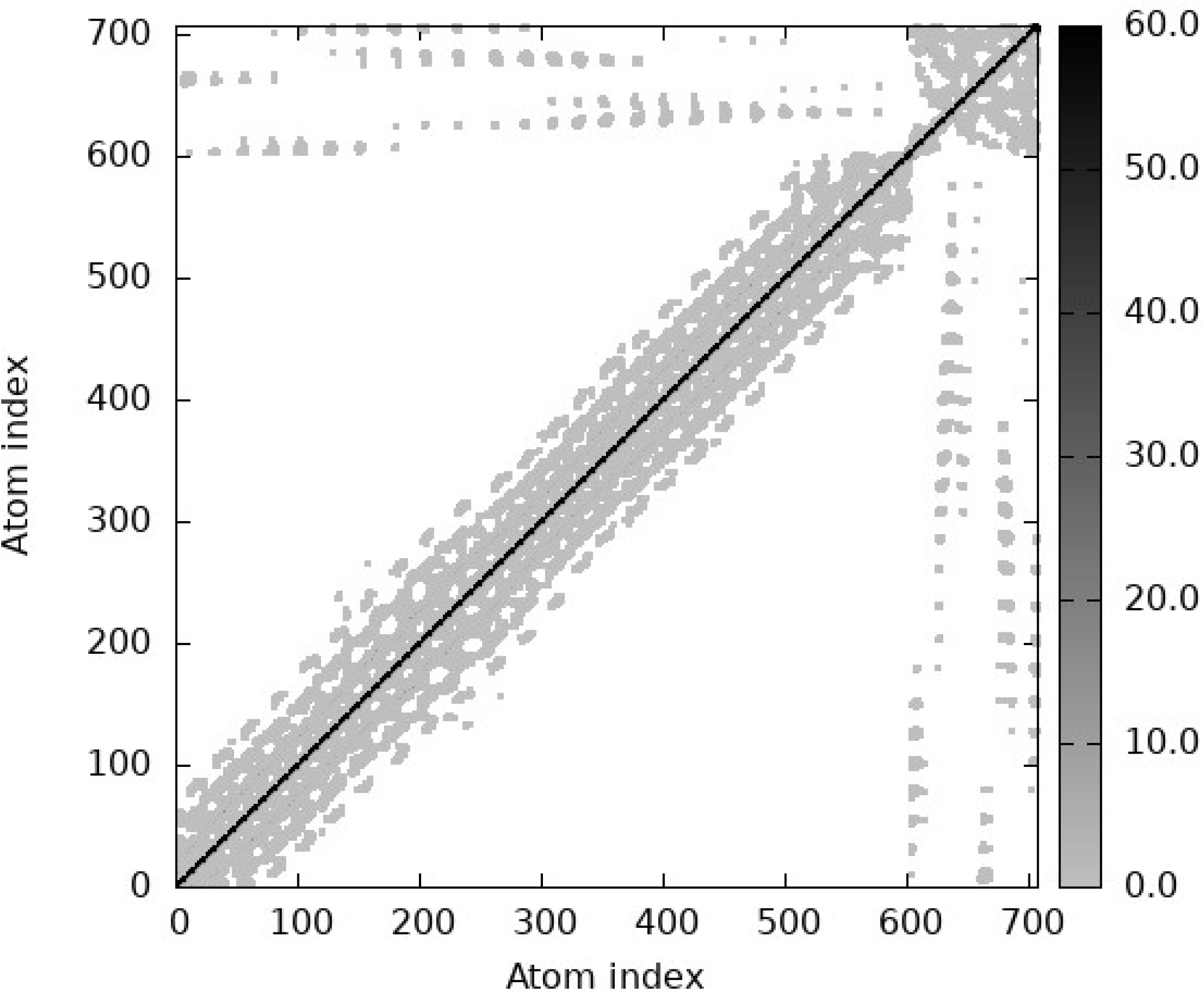

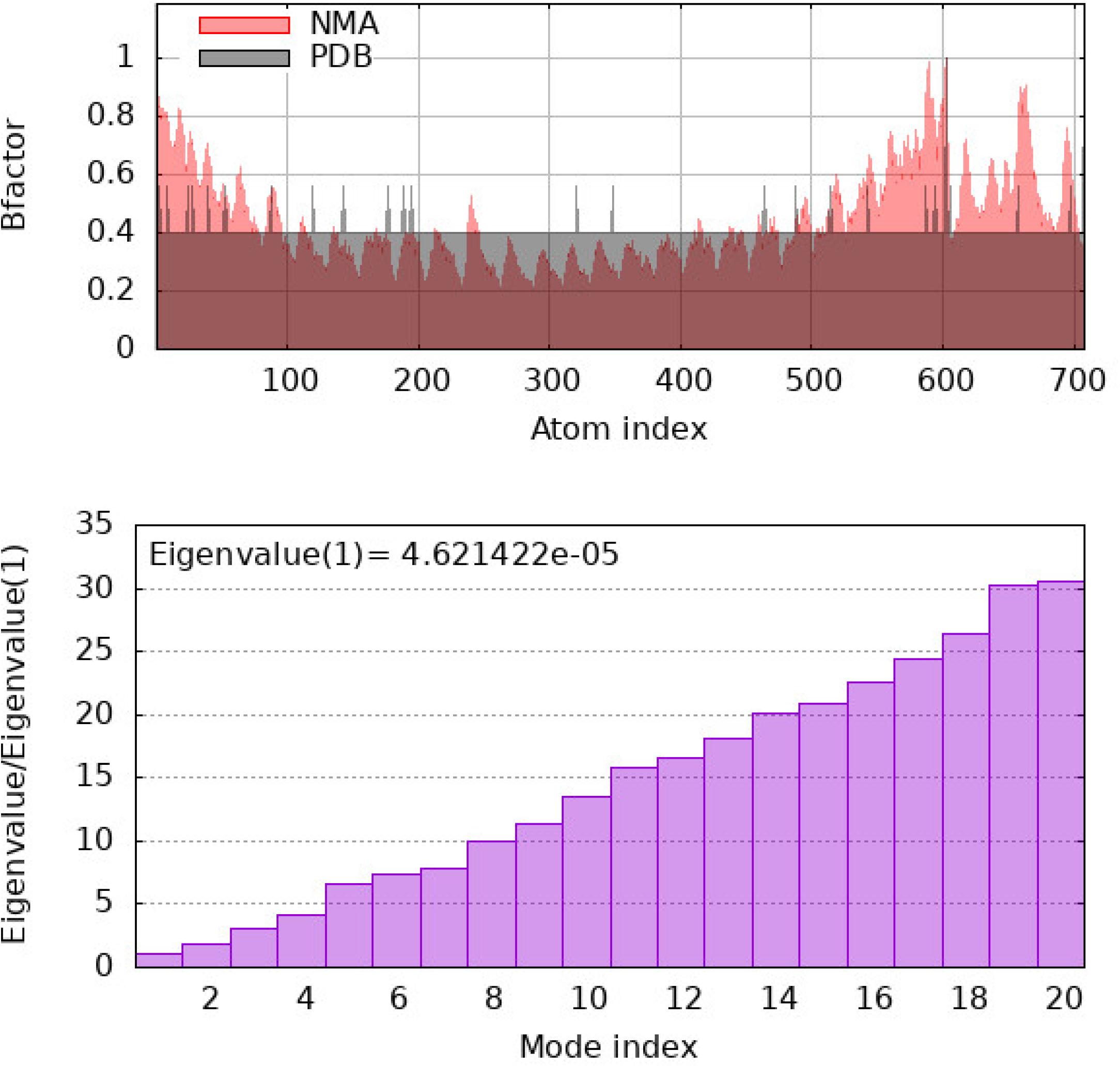

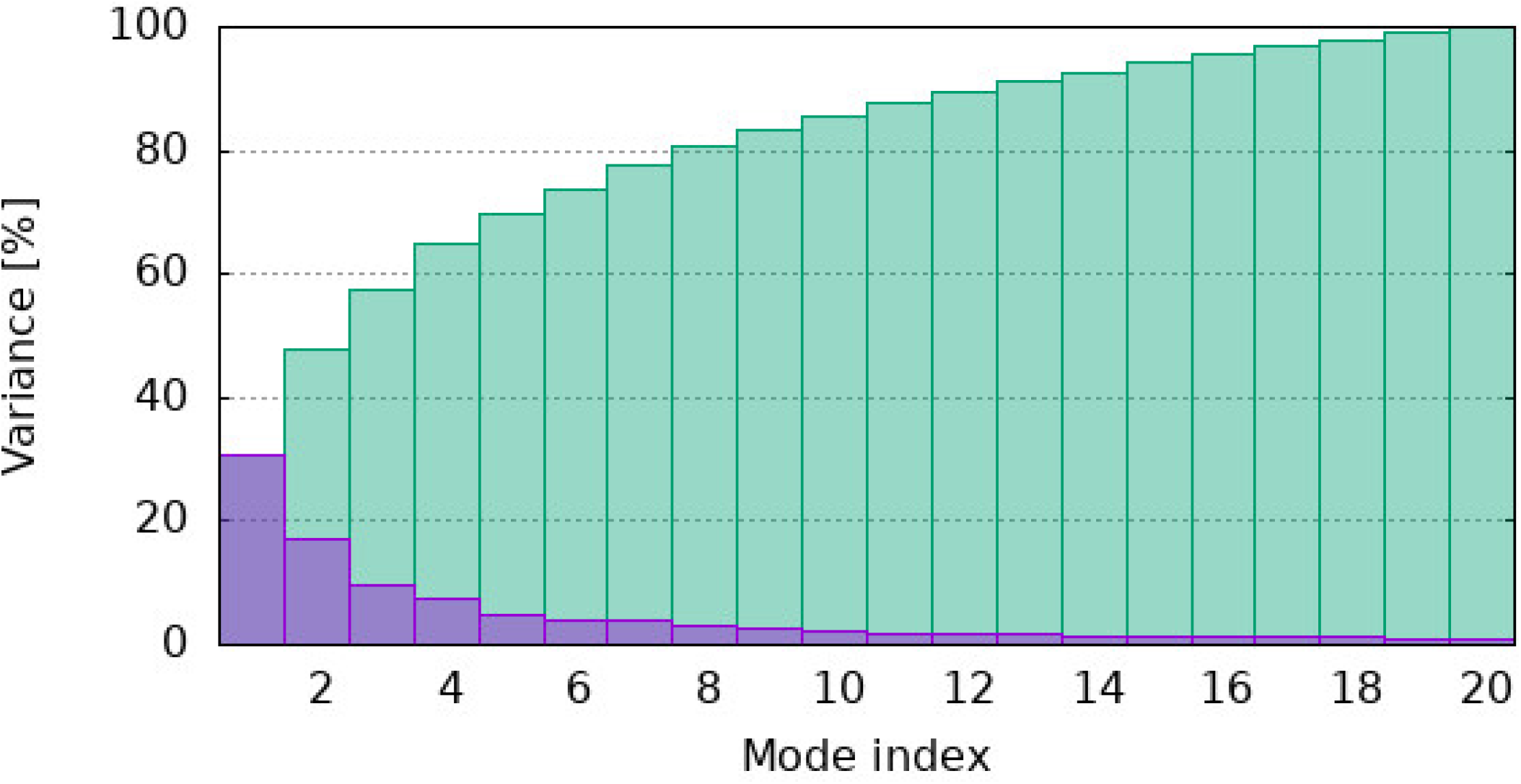

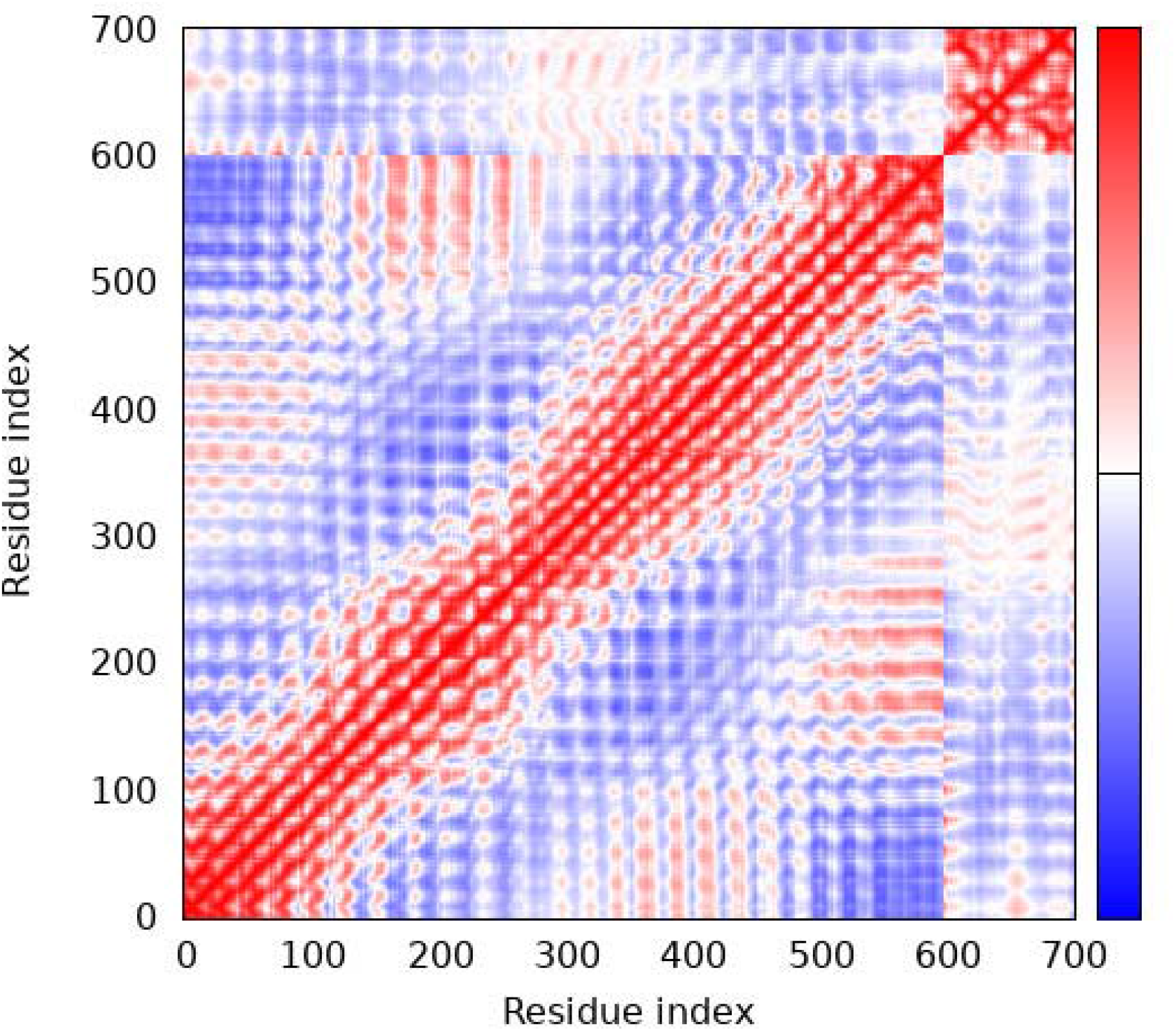

### 3.7. Computational Immune Dynamics

The C-ImmSim agent-based simulation provided a comprehensive predictive map of the vaccine’s clinical efficacy over a 70-day prime-boost-boost regimen (Rapin et al., 2010):

- **Humoral Immunity:** The initial simulated injection resulted in rapid antigen clearance, temporally correlated with a massive primary spike in Immunoglobulin M (IgM) titers. Subsequent booster doses at days 28 and 56 triggered successful B-cell class-switch recombination, manifesting as rapid, sustained, and exponentially higher expansions of IgG1 and IgG2 antibody titers, establishing long-lived humoral memory.
- **Cellular Immunity:** The cellular response was marked by a rapid expansion of active CD4+ T-helper cells, peaking around day 15 following the initial dose. This established the helper memory populations required to orchestrate comprehensive adaptive immunity.
- **Cytokine Profile:** Cytokine signaling assays revealed elevated, intense, and sustained concentrations of Interferon-gamma (IFN-γ) and Interleukin-2 (IL-2). This specific biochemical signature confirms the induction of a potent, Th1-polarized inflammatory response.

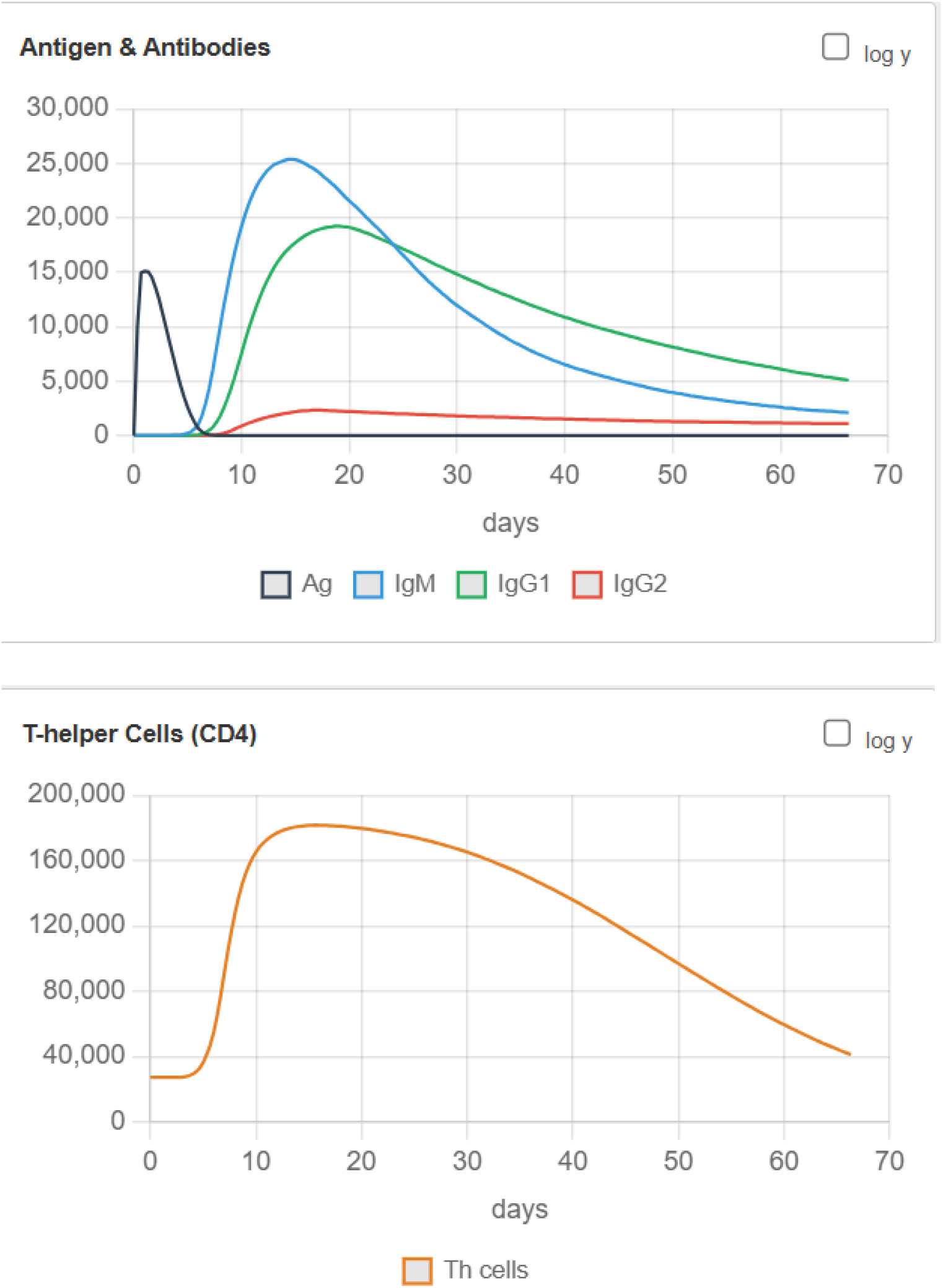

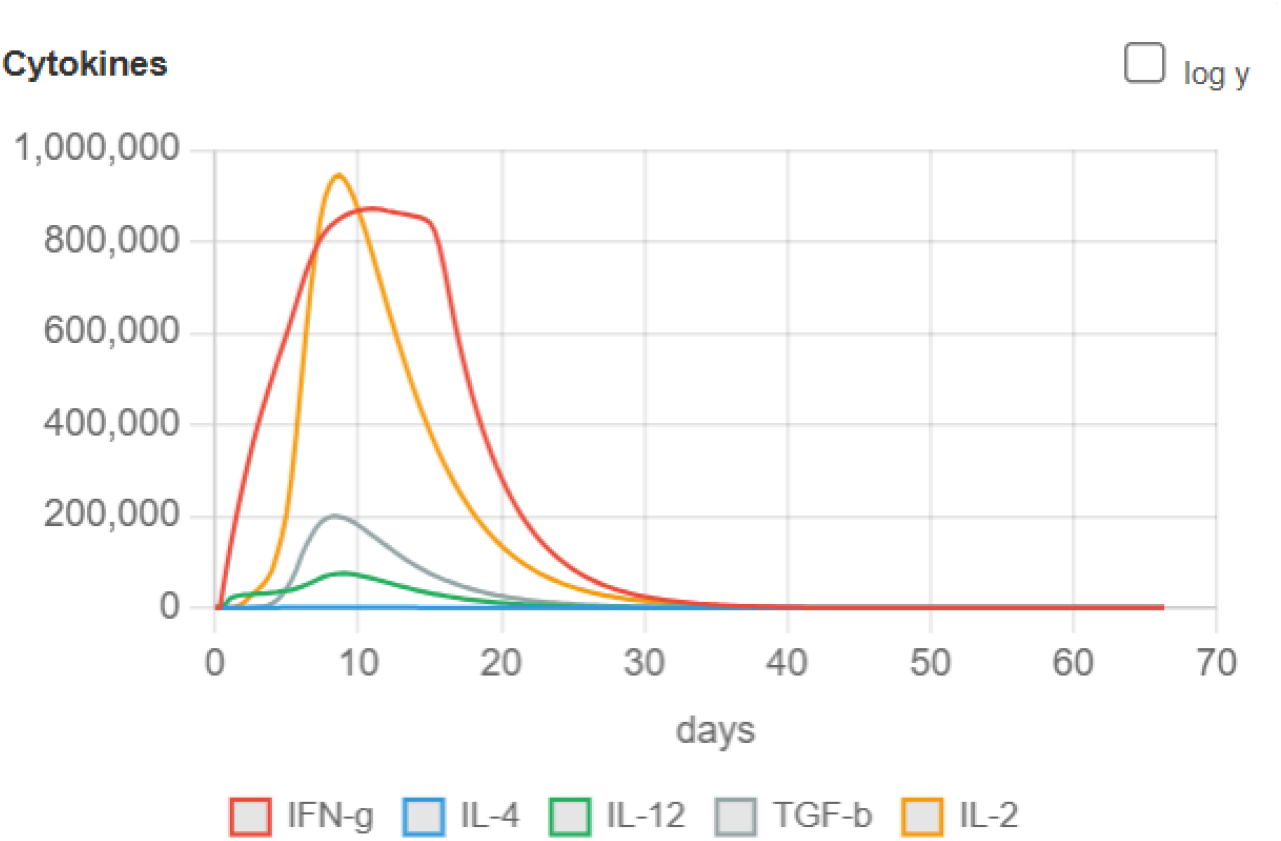

## 4. Discussion

The escalating threat of arthropod-borne viruses that cause encephalitis is a major global health concern. The Chandipura virus (CHPV) is particularly alarming due to its rapid neuro-invasive pathogenesis, which severely limits the efficacy of post-exposure treatments. Consequently, the development of a robust prophylactic vaccine has become a critical public health priority. This study presents comprehensive data detailing the design and evaluation of a 104-amino-acid recombinant construct using a rigorous in silico methodology.

One of the major limitations in multi-epitope vaccine design is the accidental inclusion of epitopes that induce IgE hypersensitivity or intrinsic cell cytotoxicity. The analysis of our construct with AllerTOP (Dimitrov et al., 2014) showed that the selected sequence does not share structural homology with any known allergens. Such dissimilarity significantly reduces the risk of adverse effects. This, together with the non-toxic characteristic confirmed by ToxinPred (Gupta et al., 2013), establishes a robust biological safety profile for our proposed vaccine candidate.

From a manufacturing point of view, the vaccine has great qualities. The codon optimization metrics—the CAI of 0.956 and GC content of 61.48%—ensure that the engineered sequence can be made at a low cost and produced in large amounts in common bacterial systems, avoiding the typical translation problems that happen when wild-type viral genomes are expressed (Grote et al., 2005). Also, the vaccine was designed to solve the problems that often cause issues with experimental peptide vaccines, like not dissolving and breaking down quickly. ExPASy ProtParam shows that the vaccine’s negative GRAVY score of –0.350 makes sure it stays dissolved in the body, while the Instability Index of 39.92 means the vaccine is stable enough to be stored and shipped (Gasteiger et al., 2005). In addition, the IEDB population coverage analysis shows that the vaccine’s combined epitopes cover more than 94% of the world and almost 90% of South Asia, going beyond the genetic limits set by HLA polymorphism and proving that the vaccine can be used widely in highly affected Indian groups. Immunologically, the success of any vaccine depends on its ability to be recognized by host antigen-presenting cells. Toll-like Receptor 4 serves as a watcher of the innate immune system responsible for recognizing viral surface glycoproteins. Our HawkDock MM/GBSA thermodynamic analysis shows mathematically that the vaccine peptide has the shape needed to bind tightly to TLR4, driven by a very good free energy change of -29.12 kcal/mol (Weng et al., 2019).

However, static thermodynamics only give a snapshot of molecular affinity. In a living body, proteins move constantly. By using Normal Mode Analysis through the iMODS server, this study clearly showed the structural stability of the vaccine-receptor complex (López-Blanco et al., 2014). The low eigenvalue of 4.621422×10-5 proves that the vaccine-receptor complex has great motion stiffness, needing almost no external energy to keep its docked shape. The covariance matrix shows correlated residue motions, and the elastic network shows stiff connections. These facts of stability guarantee that the vaccine will stay attached to the TLR4 receptor long enough to activate the MyD88-dependent signaling cascade. This activation leads to the NF-κB transcription factor turning on and the subsequent maturation of the APC.

This stable molecular trigger was clearly shown in the agent-based C-ImmSim simulation results (Rapin et al., 2010). The vaccine produced high, sustained IgG1 and IgG2 titers, which shows that the vaccine can give long-lasting neutralizing immunity against circulating virions. For a virus like CHPV that quickly enters the nervous system, the simulation also showed that the vaccine construct causes a large burst of IFN-γ. High amounts of IFN-γ in the body are essential to turn on macrophages and cytotoxic CD8+ T-cells, which helps the vaccine to focus a Th1-polarized inflammatory reaction that is needed to clear viral pathogens that have already entered target host cells.

This study carried out a computational design and full evaluation of a new multi-epitope vaccine for the Chandipura virus. The 104-amino-acid peptide we built was safe, showing no allergy or toxicity. It was stable in physicochemical tests, making it suitable for an industrial drug. The peptide matched codon usage, making it easy to produce in large amounts, and it bound strongly to the human TLR4 receptor. We added Normal Mode Analysis to confirm that the multi-epitope vaccine keeps its shape when the receptor moves like it does inside the body. Agent-based immune simulations predicted a Th1-type response, both in antibodies and in cells. Together, this multi-epitope vaccine looks very promising. These strong computational results give a scientific basis to move quickly to make the peptide in the lab, test it in cells, and then do animal studies.

## Acknowledgements

The author wishes to express profound gratitude to Dr. Himanshu Singh, Head of the Department of Biomedical Engineering, for his invaluable guidance, mentorship, and continuous encouragement throughout the conception and execution of this immunoinformatics project. The author also extends their appreciation to the Department of Biotechnology at Lovely Professional University for providing the academic framework and computational resources necessary to complete this research.

## Declaration of Generative AI and AI-Assisted Technologies

*During the preparation of this work, the author(s) used Google Gemini in order to polish the English language and correct grammatical errors. After using this tool, the author(s) reviewed and edited the content as needed and take full responsibility for the content of the publication*.

## References

Dimitrov, I., Bangov, I., Flower, D. R., & Doytchinova, I. (2014). AllerTOP v.2—a server for in silico prediction of allergens. Journal of Molecular Modeling, 20(6), 2278. 10.1007/s00894-014-2278-5

Doytchinova, I. A., & Flower, D. R. (2007). VaxiJen: a server for prediction of protective antigens, tumour antigens and subunit vaccines. BMC Bioinformatics, 8, 4. 10.1186/1471-2105-8-4

Gasteiger, E., Hoogland, C., Gattiker, A., Duvaud, S., Wilkins, M. R., Appel, R. D., & Bairoch, A. (2005). Protein Identification and Analysis Tools on the ExPASy Server. In J. M. Walker (Ed.), The Proteomics Protocols Handbook (pp. 571–607). Humana Press. 10.1385/1-59259-890-0:571

Grote, A., Hiller, K., Scheer, M., Munch, R., Nortemann, B., Hempel, D. C., & Jahn, D. (2005). JCat: a novel tool to adapt codon usage of a target gene to its potential expression host. Nucleic Acids Research, 33, W526–W531. 10.1093/nar/gki376

Gupta, S., Kapoor, P., Chaudhary, K., Gautam, A., Kumar, R., & Raghava, G. P. S. (2013). In Silico Approach for Predicting Toxicity of Peptides and Proteins. PLoS ONE, 8(9), e73957. 10.1371/journal.pone.0073957

López-Blanco, J. R., Aliaga, J. I., Quintana-Ortí, E. S., & Chacón, P. (2014). iMODS: internal coordinates normal mode analysis server. Nucleic Acids Research, 42, W271–W276. 10.1093/nar/gku339

Rapin, N., Lund, O., Bernaschi, M., & Castiglione, F. (2010). Computational Immunology Meets Bioinformatics: The Use of Prediction Tools for Molecular Binding in the Simulation of the Immune System. PLoS ONE, 5(4), e9862. 10.1371/journal.pone.0009862

Weng, G., Wang, E., Wang, Z., Liu, H., Zhu, F., Li, D., & Hou, T. (2019). HawkDock: a web server to predict and analyze the protein–protein complex based on computational docking and MM/GBSA. Nucleic Acids Research, 47, W322–W330. 10.1093/nar/gkz397

